# Next-generation diarylquinolines improve sterilizing activity of regimens with pretomanid and the novel oxazolidinone TBI-223 in a mouse tuberculosis model

**DOI:** 10.1101/2023.01.10.523519

**Authors:** Si-Yang Li, Paul J. Converse, Fabrice Betoudji, Jin Lee, Khisimuzi Mdluli, Anna Upton, Nader Fotouhi, Eric L. Nuermberger

## Abstract

A regimen comprised of bedaquiline, pretomanid and linezolid (BPaL) is the first oral 6-month regimen approved by the US Food and Drug Administration and recommended by the World Health Organization for treatment of extensively drug-resistant tuberculosis. We used a well-established BALB/c mouse model of tuberculosis to evaluate the treatment-shortening potential of replacing bedaquiline with either of two new, more potent diarylquinolines in early clinical trials, TBAJ-587 and TBAJ-876. We also evaluated the effect of replacing linezolid with a new oxazolidinone, TBI-223, exhibiting a larger safety margin with respect to mitochondrial toxicity in preclinical studies. Replacing bedaquiline with TBAJ-587 at the same 25 mg/kg dose significantly reduced the proportion of mice relapsing after 2 months of treatment, while replacing linezolid with TBI-223 at the same 100 mg/kg dose did not significantly change the proportion of mice relapsing. Replacing linezolid or TBI-223 with sutezolid in combination with TBAJ-587 and pretomanid significantly reduced the proportion of mice relapsing. In combination with pretomanid and TBI-223, TBAJ-876 at 6.25 mg/kg was equipotent to TBAJ-587 at 25 mg/kg. We conclude that replacement of bedaquiline with these more efficacious and potentially safer diarylquinolines and replacement of linezolid with potentially safer and at least as efficacious oxazolidinones in the clinically successful BPaL regimen may lead to superior regimens capable of treating both drug-susceptible and drug-resistant TB more effectively and safely.

---

Bedaquiline (B) has transformed the treatment of multidrug- and extensively drug-resistant tuberculosis (MDR/XDR-TB). For example, the novel regimen comprised of bedaquiline, pretomanid (Pa) and linezolid (L), abbreviated as BPaL, has proven efficacy as a 6-month oral regimen to treat MDR/XDR-TB and is the first and only such regimen approved for this indication (1, 2). However, in order to achieve the oft-stated objective of even shorter treatment regimens appropriate for both drug-susceptible TB and MDR/XDR-TB and to more effectively counter the threat of emerging bedaquiline resistance, further improvements in the BPaL regimen will be required. Shorter regimens may be achieved with inclusion of more potent drugs. Indeed, two next-generation diarylquinoline (DARQ) drugs with more potent activity than bedaquiline are now in phase 1 clinical trials, TBAJ-587 (S587) and TBAJ-876 (S876), (ClinicalTrials.gov identifiers: NCT04890535 and NCT04493671, respectively). We recently used a well-established BALB/c mouse infection model of TB to demonstrate the superior bactericidal activity of novel regimens in which either new DARQ is used in place of bedaquiline in the BPaL regimen (3, 4). The newer DARQs also retained greater activity against isogenic strains with reduced bedaquiline susceptibility due to mutations in the *mmpL5/mmpS5* repressor, *Rv0678* (also known as *mmpR5*). Despite these promising observations of the superior bactericidal activity of these newer DARQs, evaluation of sterilizing activity using the endpoint of relapse-free cure in this mouse model is considered a more reliable indication of the treatment-shortening potential of a new regimen (5, 6).

Despite its demonstrated efficacy as a short-course oral regimen, the clinical use of BPaL carries significant safety concerns related to the hematologic and neurologic toxicity of linezolid (1, 2). A safer oxazolidinone could reduce the need for safety monitoring, dose reductions and drug holidays and perhaps expand the utility of a DARQ+pretomanid+oxazolidinone regimen to the treatment of drug-susceptible TB. TBI-223 (O) is a new oxazolidinone with *in vitro* potency against *Mycobacterium tuberculosis* approaching that of linezolid that has demonstrated a much lower risk of mitochondrial toxicity in preclinical safety studies. It is currently being evaluated in a phase 1 multiple ascending dose study (ClinicalTrials.gov identifier: NCT03758612). Sutezolid (U) is another oxazolidinone, now in a phase 2b trial (ClinicalTrials.gov Identifier: NCT03959566) that has more potent activity than linezolid in mouse models and may also have reduced mitochondrial toxicity (7–9). Hence, these newer oxazolidinones warrant further evaluation as replacements for linezolid in combinations with a DARQ and pretomanid.

In this study, we evaluated the sterilizing activity of S587PaL and S876PaL in comparison to BPaL and assessed whether other potentially safer, oxazolidinones, TBI-223 and sutezolid, can meet or exceed the sterilizing activity of linezolid in combination with a DARQ and pretomanid in the mouse model in which the sterilizing activity of the BPaL regimen was first demonstrated (7).

## RESULTS

### Pharmacokinetics of the diarylquinolines in mice

Plasma pharmacokinetics (PK) profiles were determined after 1 and 7 consecutive days of dosing in uninfected BALB/c mice. Plasma PK parameters of S587, S876, and bedaquiline and their active metabolites at different doses are shown in Table 1.

**Table 1.** Plasma PK parameter values for the three diarylquinolines under study.

| Parameter | Drug |  |  |  |  |
| --- | --- | --- | --- | --- | --- |
|  | S587 |  | S876 |  | Bedaquiline |
| Dose (mg/kg) | 12.5 | 25 | 3.125 | 6.25 | 25 |
| $T_{1/2}$* Day 1 | 10.1±2.1 | 9.09±1.6 | 8.99±1.36 | 8.42±0.96 | 6.63±1.05 |
| Day 7 | 55.3±8.0 | 63.4±4.5 | 69±30.6 | 56.8±2.0 | 112±56 |
| $T_{max}$ (hr)* Day 1 | 1±0 | 1±0 | 1±0 | 1±0 | 1.67±0.58 |
| Day 7 | 1.33±0.58 | 1.33±0.58 | 1±0 | 1±0 | 1.67±0.58 |
| $C_{max}$ (ng/ml)* Day 1 | 1024±175 | 2173±195 | 220±110 | 413±85 | 2867±1305 |
| Day 7 | 1533±352 | 2813±865 | 182±43 | 468±147 | 1250±70 |
| AUC<sub>0-24h</sub> (hr/ng/ml)* |  |  |  |  |  |
| Day 1 | 8203±1662 | 16214±2260 | 1260±165 | 2545±592 | 21151±8471 |
| Day 7 | 13052±3415 | 28841±2987 | 1470±372 | 4030±1234 | 9628±1418 |
| Metabolite |  |  |  |  |  |
|  | S587-M3 |  | S876-M3 |  | Bedaquiline-M2 |
| $T_{1/2}$* Day 1 | 29.9±7.3 | 37.3±5.4 | NA | NA | 34.3±6.1 |
| Day 7 | 50.9±10.7 | 53.6±8 | 32.2±8.4 | 30.3±4 | 50.7±6.4 |
| $T_{max}$ (hr)* Day 1 | 8±0 | 8±0 | 6.67±2.31 | 6±3.46 | 12±10.6 |
| Day 7 | 4.67±3.06 | 5.33±2.31 | 3.33±1.15 | 6±3.46 | 4.67±3.06 |
| $C_{max}$ (ng/ml)* Day 1 | 563±65 | 1051±94 | 181±14 | 318±50 | 809±197 |
| Day 7 | 1350±113 | 2107±142 | 330±43 | 760±131 | 2677±124 |
| AUC<sub>0-24h</sub> (hr/ng/ml)* |  |  |  |  |  |
| Day 1 | 11073±1301 | 20797±1589 | 3365±138 | 6076±492 | 14940±3632 |
| Day 7 | 24922±1075 | 41557±4339 | 6027±328 | 15564±2227 | 55130±3049 |
| *values are mean ± sd |  |  |  |  |  |

#### Experiment 1

The dose-ranging activity of S587 at doses of 5, 10, 25, 50 and 100 mg/kg in combination with PaL was evaluated in a sub-acute, high-dose aerosol infection model of TB in BALB/c mice. After 2 weeks of treatment, as shown in Fig 1A, a S587 dose-dependent reduction in lung CFU was observed for doses up to 25 mg/kg with no further increase in activity with further dose increases to 50 and 100 mg/kg. PaL was less active than all S587PaL regimens (p<0.0001), except for S587_5_PaL. S587_25_PaL was more active than S587_5_PaL (p<0.0001) and S587_10_PaL (p=0.0031) but not different from S587_50_PaL or S587_100_PaL. After 4 weeks of treatment, a S587 dose-dependent response was again observed up to 25 mg/kg, whereas the 25 mg/kg dose was not significantly different from the 50 mg/kg dose and was more active than the 100 mg/kg dose (p=0.0456) (Table S1). All S587PaL regimens and BPaL were more active than PaL (p<0.0001) at this time point. S587_25_PaL was again significantly more active than S587_5_PaL and S587_10_PaL (p<0.0001). BPaL was less active than S587PaL when S587 doses were ≥25 mg/kg (p<0.0001, <0.0001, and 0.0010, in order of ascending S587 dose) but more active than S587_5_PaL.

**FIG 1.**
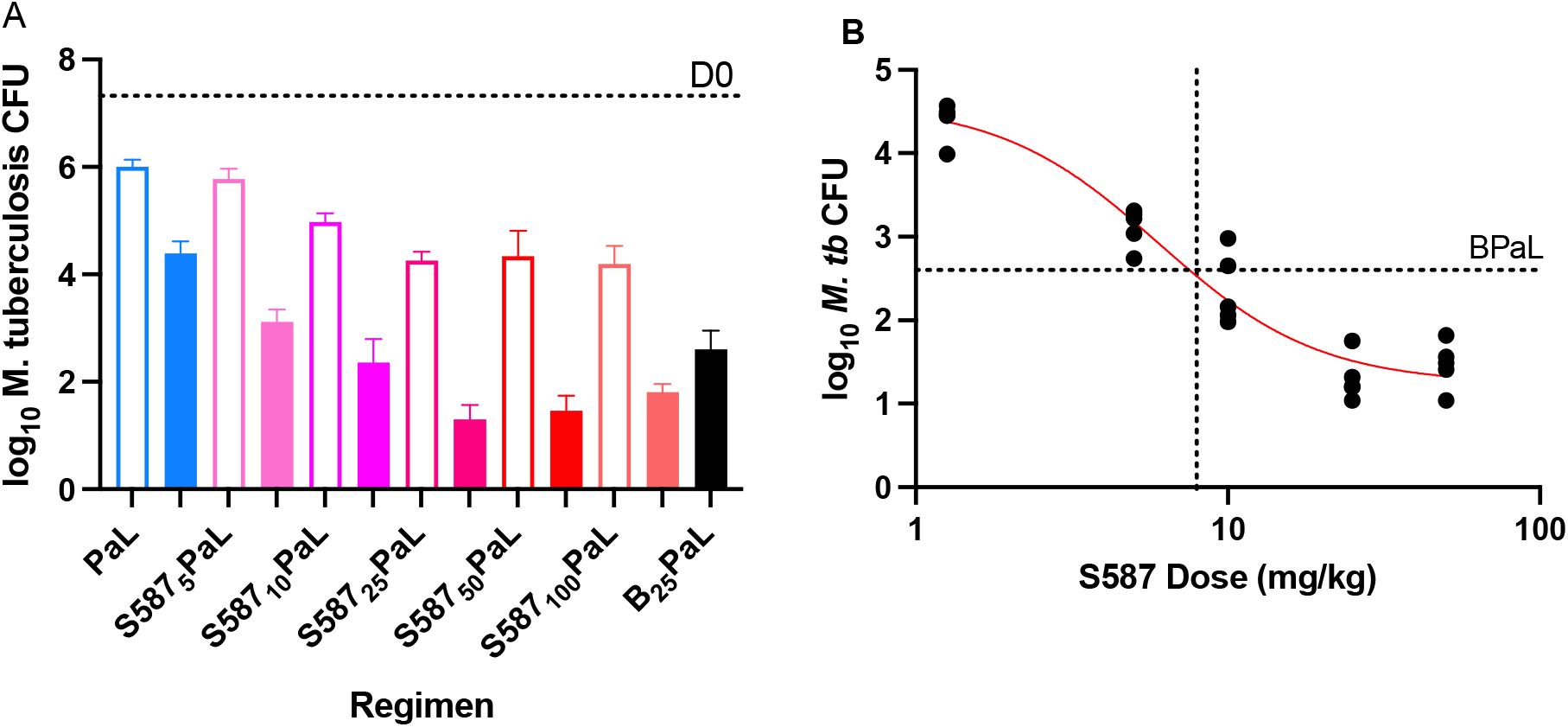
Dose-ranging activity of S587 combined with PaL. A) CFU results after 2 weeks (open bars) and 4 weeks of treatment (solid bars) are shown. S587-containing regimens are in red shades, BPaL in black, and PaL alone in blue. B) A sigmoidal dose-response curve fit to the 4-week treatment results was used to estimate the S587 dose equivalent to BDQ 25 mg/kg in combination with PaL. The latter regimen after 4 weeks of treatment reduced the burden of *M. tuberculosis* to 2.53±0.32 CFU as indicated by the horizontal line marked BPaL. The S587 dose (7.6 mg/kg) in the SPaL regimen that would achieve such a reduction is indicated by the vertical dotted line

Nonlinear regression analysis was used to fit a sigmoidal Em_ax_ curve to describe the dose-response relationship for the contribution of S587 to the S587PaL regimen. By interpolation, a S587 dose of 7.6 mg/kg was estimated to be equivalent to bedaquiline at 25 mg/kg when added to PaL (Fig. 1B). The ED_90_ for S587 was estimated to be 22.6 mg/kg. Thus, S587 added bactericidal activity to PaL, even at the lowest dose (5 mg/kg) tested. At 7.6 mg/kg S587 was equivalent to bedaquiline at 25 mg/kg while S587 at 25 mg/kg was superior to bedaquiline at 25 mg/kg, as reported previously by Xu et al. (4).

#### Experiment 2

After S587 was confirmed to have more potent bactericidal activity than bedaquiline, a relapse study was conducted to assess the treatment-shortening potential of replacing bedaquiline with S587 at the same 25 mg/kg dose in the BPaL regimen. BPaL plus the well-known sterilizing drug pyrazinamide (Z) was included as a comparator. After four weeks of treatment the mean lung CFU count was lower in mice receiving S587PaL (p=0.0548) or BPaLZ (p<0.0001), compared to BPaL. Mice were treated for an additional two (W6) or four weeks (W8) and then left untreated for an additional 12 weeks. At the W6 (+12) relapse time point, all 15 BPaL-treated mice relapsed but only 10 of 14 S 587_25_PaL-treated mice relapsed (p=0.0421). At the W8 (+12) relapse time point, all 15 BPaL-treated mice again relapsed but only 4 of 13 S587_25_PaL-treated mice relapsed (p<0.0001) (Table 2). None of the BPaLZ-treated mice relapsed at either time point. Thus, S587_25_PaL has greater bactericidal activity and sterilizing activity compared to BPaL, but replacing bedaquiline with S587 does not increase the sterilizing activity as much as adding pyrazinamide.

**Table 2.** Lung CFU counts assessed during treatment and proportion of mice relapsing after treatment completion in experiment 2

| Regimen | Mean lung log<sub>10</sub> CFU count (± SD) | | | Proportion relapsing after treatment for: | |
| --- | --- | --- | --- | --- | --- |
|  | W-2 | D0 | W4 | W6 (+12) | W8 (+12) |
| Untreated | 3.98±0.13 | 7.17±0.10 | | | |
| B<sub>25</sub>Pa<sub>100</sub>L<sub>100</sub> |  |  | 2.85±0.42 | 15/15 | 15/15 |
| BPaLZ<sub>150</sub> |  |  | 0.46±0.26 | 0/15 | 0/15 |
| S587<sub>25</sub>Pa<sub>100</sub>L<sub>100</sub> |  |  | 2.03±0.63 | 10/14* | 4/13** |

W-2=1 day after aerosol infection (n= mice); D0= day of treatment initiation, 2 weeks after infection. W4=treated for 4 weeks; W6+(12) = treated for 6 weeks, held for an additional 12 weeks without treatment, then sacrificed to determine the proportion with relapse, etc. From D0, 15 mice were allocated for relapse assessment at each time point indicated by proportions in the table. Due to an unscheduled death due to unknown causes 4 weeks after completing 6 weeks of treatment with S587PaL (\*), only 14 mice were assessable in this arm; cultures of 2 mouse lungs (\*\*) were contaminated and could not be assessed.

#### Experiment 3

A follow-up relapse study was conducted to confirm the superior sterilizing activity obtained by substituting S587 at 25 mg/kg for bedaquiline in the BPaL regimen and also to assess the potential of TBI-223 to replace linezolid. The geometric mean MICs of linezolid and TBI-223 were 1 and 3.175 μg/ml, respectively, against the infecting strain. After 4 weeks of treatment (W4), the bactericidal activity of S587 in combination with PaL was again significantly (p=0.0008) greater than BPaL. BPa was significantly less active than BPaL (p=0.0019) and BPaO (p=0.0299), indicating that both oxazolidinones added bactericidal activity to the combination (Table 3). The difference between BPaL and BPaO was not statistically significant. After 8 weeks of treatment (W8), BPa was again significantly less active than BPaL (p=0.001) and BPaO (p=0.0153). The proportion of mice relapsing after 8 weeks of treatment with BPaL followed by 12 weeks without treatment was significantly higher than the proportion of mice relapsing after either 6 (p=0.0352) or 8 weeks (p<0.0001) of S587PaL. Likewise, the proportion of mice relapsing after 12 weeks of treatment with BPaL was significantly higher than the proportion of mice relapsing after either 8 (p=0.0352) or 12 weeks (p=0.0063) of S587PaL. At W12 (+12) nearly all mice treated with BPa alone relapsed, while significantly fewer mice treated with BPaL (p=0.0142) or BPaO (p<0.0001) relapsed. The difference between BPaL and BPaO was not statistically significant (p=0.1086). Thus, both oxazolidinones contributed similar bactericidal and sterilizing activity to the BPa combination. Substituting S587 for bedaquiline in the combination with PaL enhanced both bactericidal and sterilizing activity, shortening the treatment duration needed to prevent relapse in half of the mice by approximately 6 weeks.

**Table 3.** Lung CFU counts assessed during treatment and proportion of mice relapsing after treatment completion in experiment 3

| Regimen | Mean lung log<sub>10</sub> CFU count (±SD) |  |  |  | Proportion relapsing after treatment for: |  |  |
| --- | --- | --- | --- | --- | --- | --- | --- |
|  | W-2 | D0 | W4 | W8 | W6 (+12) | W8 (+12) | W12 (+12) |
| Untreated | 4.66±0.08 | 8.85±0.15 |  |  |  |  |  |
| B<sub>25</sub>Pa<sub>100</sub> |  |  | 5.97±0.34 | 2.96±0.42 |  |  | 14/15 |
| B<sub>25</sub>Pa<sub>100</sub>L<sub>100</sub> |  |  | 4.48±0.41 | 1.00±0.90<sup>1</sup> |  | 14/15 | 7/15 |
| S587<sub>25</sub>Pa<sub>100</sub>L<sub>100</sub> |  |  | 2.82±0.37 | 0.00±0.00 | 8/15 | 1/15 | 0/15 |
| B<sub>25</sub>Pa<sub>100</sub>O<sub>100</sub> |  |  | 4.92±0.40 | 1.62±0.54 |  | 15/15 | 2/15 |

W-2=1 day after aerosol infection (n= mice); D0= day of treatment initiation, 2 weeks after infection. W4=treated for 4 weeks; W8=treated for 8 weeks; W6+(12) = treated for 6 weeks, held for an additional 12 weeks without treatment, then sacrificed to determine the proportion with relapse, etc. From D0, 15 mice were allocated for relapse assessment at each time point indicated by proportions in the table. During the 2<sup>nd</sup> week of treatment, one BPa and one BPaO mouse died due to apparent gavage accident. <sup>1</sup>One mouse was culture negative.

#### Experiment 4

A follow-up experiment was conducted to test whether TBI-223 could replace linezolid in combination with S587 and pretomanid. Sutezolid was included as an additional oxazolidinone comparator. After 4 weeks of treatment, compared to BPaL, S587_25_PaL (p=0.0021) and S587_25_PaU (p<0.0001) had superior bactericidal activity, while the difference with S587_25_PaO did not reach statistical significance (p=0.119) (Table 4). Compared to S587Pa alone, S587PaL, S587PaU, and S587PaO were statistically superior (p<0.0001, p<0.0001, p=0.0098). Only S587PaU treatment was associated with a significant reduction in the proportion of relapses after 4 weeks of treatment (p=0.0001 vs. other regimens). At W6 (+12), S587PaL and S587PaU cured all but one mouse and were significantly better than S587Pa (p=0.0001) but not different from BPaL or S587PaO (p=0.1686). Compared to S587Pa, S587PaO had significantly more sterilizing activity at this time point (p=0.0213). Thus, the benefit of replacing bedaquiline with S587 in the BPaL regimen was again confirmed in terms of bactericidal activity but not sterilizing activity because there were fewer than the expected number of relapses at W6 in the BPaL group. Furthermore, TBI-223 added bactericidal and sterilizing activity to the S587Pa backbone and resulted in a regimen at least as effective as BPaL and not significantly worse than S587PaL. This experiment also demonstrated the superior bactericidal and sterilizing activity of replacing linezolid with sutezolid in combination with S587Pa.

**Table 4.**
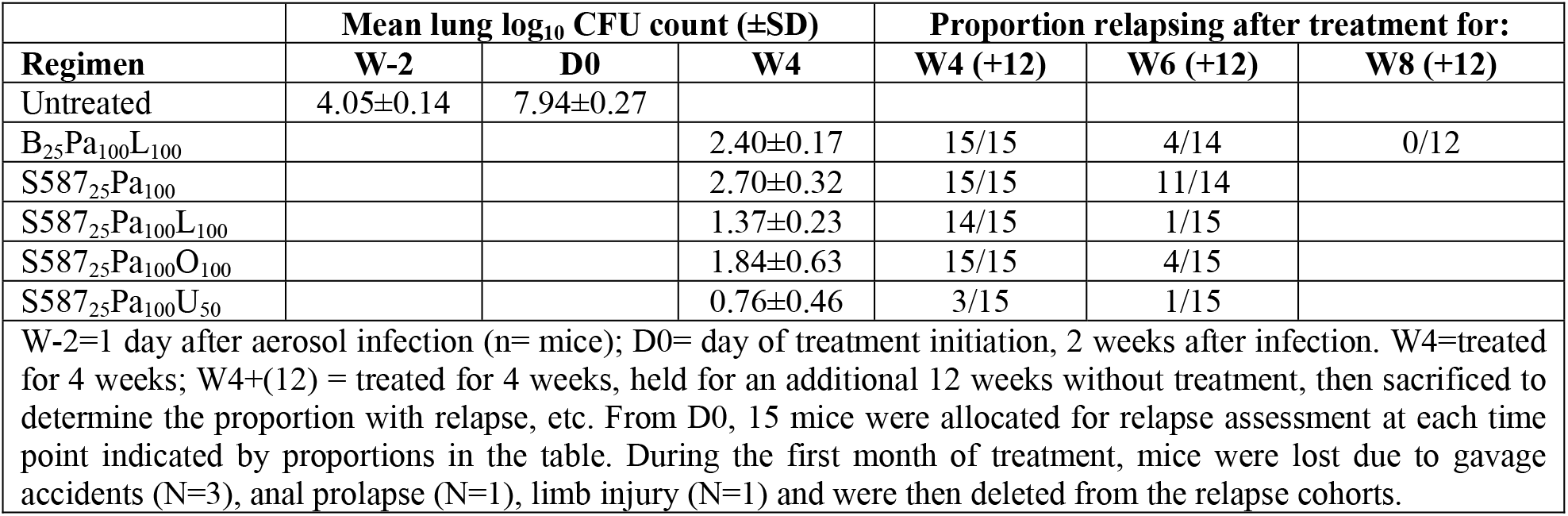
Lung CFU counts assessed during treatment and proportion of mice relapsing after treatment completion in experiment 4

#### Experiment 5

In the first head-to-head comparison of S587 and S876 in combination therapy, we compared the bactericidal activity of S876 to that of bedaquiline and S587 in combination with PaO. Based on its greater potency relative to bedaquiline and S587 observed previously, S876 was dosed at 6.25 mg/kg. After four weeks of treatment S587 showed dose-ranging (at 25 and 50 mg/kg) enhancement of bactericidal activity and both S587 and S876 were highly significantly (p<0.0001) more active than bedaquiline when administered in a regimen with PaO (Fig. 2). S876 at 6.25 mg/kg was approximately equipotent with S587 at 25 mg/kg.

**FIG 2.**
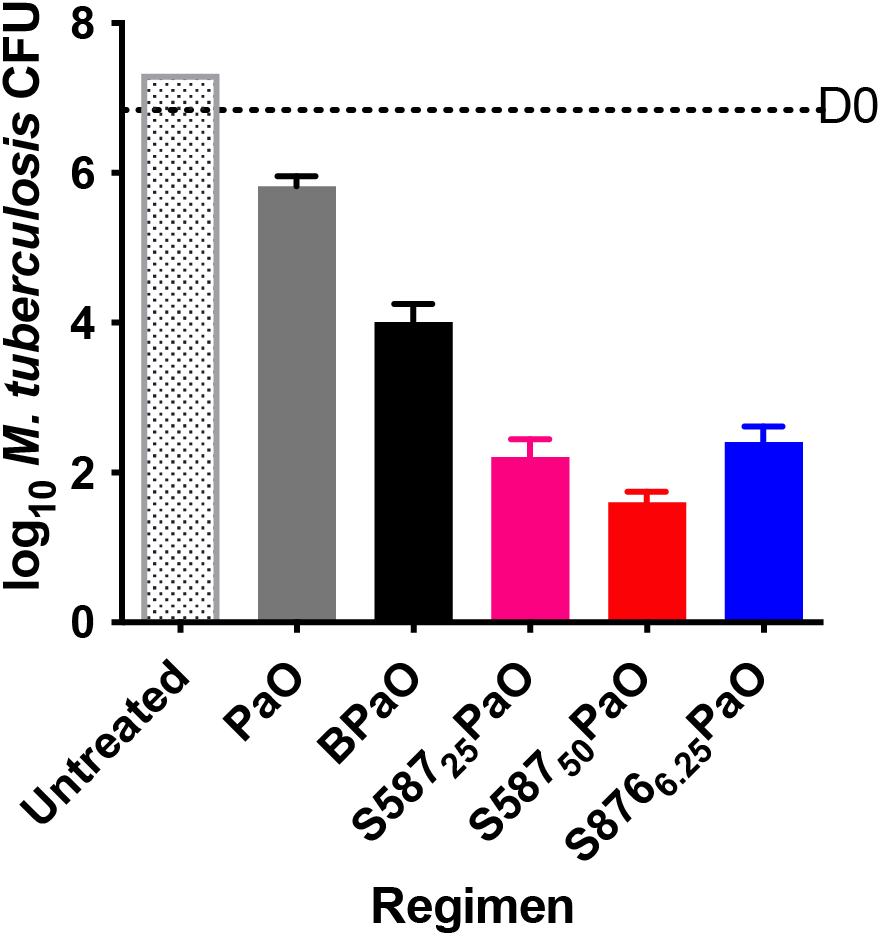
Bactericidal activity after four weeks of treatment with S587 or S876 in combination with pretomanid (Pa) and TBI-223 (O).

#### Experiment 6

The final experiment compared the dose-ranging sterilizing activity of S587 and S876 to bedaquiline when administered together with PaO. At the beginning of treatment, there were 6.58±0.22 log_10_ CFU in the lungs. Already at the Week 4 (+12) relapse time point, owing to the low bacterial burden at the start of treatment, both S587 and S876 showed sterilizing activity, especially at the higher doses (Table 5, Fig. 3A). In combination with PaO, just 4 weeks of treatment with S587 at 50 mg/kg or S876 at 12.5 mg/kg, or 6 weeks of treatment with S587 at 25 mg/kg or S876 at 6.25 mg/kg resulted in similar proportions of mice relapsing compared to 8 weeks of treatment with bedaquiline at 25 mg/kg. At Week 6 (+12), there were not only significantly (p<0.001 vs. all other groups) more relapses in the BPaO-treated mice than in any other group, but the relapses also occurred with a high number of CFU (Fig. 3B). At the Week 8 (+12) relapse point, most mice were cured regardless of regimen, but two BPaO-treated mice still had high lung CFU burdens (Fig. 3C). In conclusion, replacing bedaquiline with S587 at 50 mg/kg or S876 12.5 mg/kg halved the treatment duration required to prevent relapse. After 4 or 6 weeks of treatment, S587 was more effective than bedaquiline at the same 25 mg/kg dose, while S876 at one-fourth the dose (6.25 or 12.5 mg/kg) was as effective in preventing relapse as S587 at 25 or 50 mg/kg, respectively, and more effective than bedaquiline.

**FIG 3.**
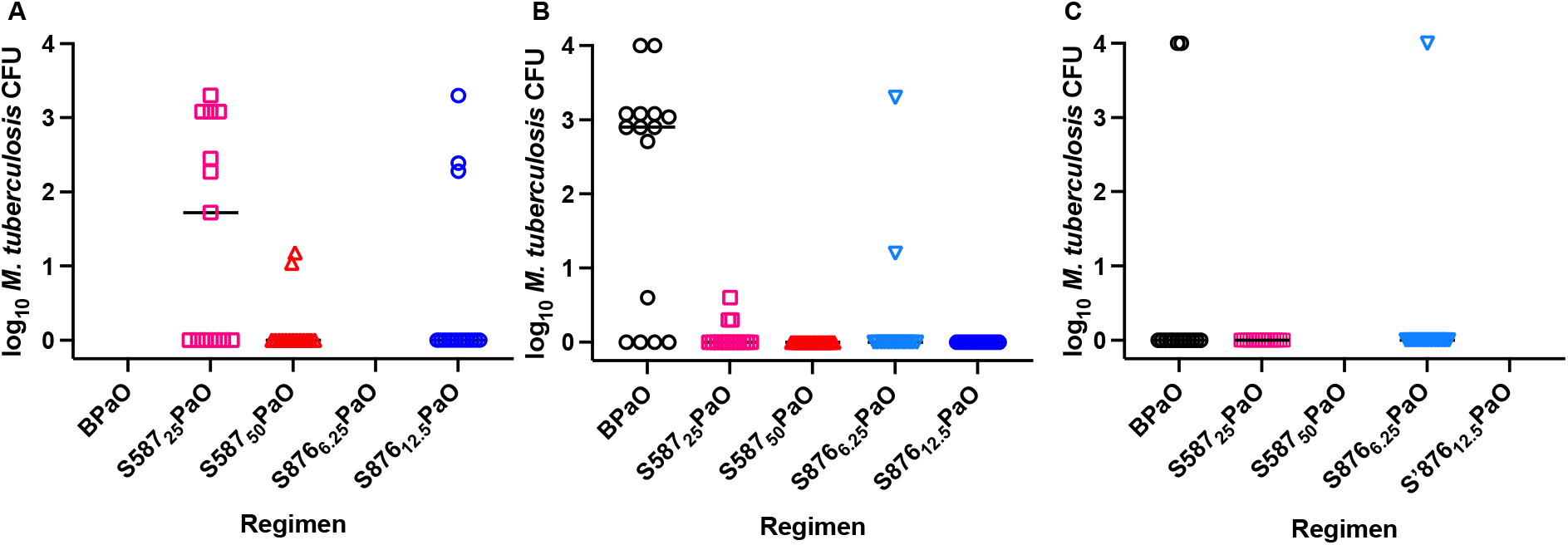
Number of CFU detected in individual mice at relapse assessment after 4 (A), 6 (B), and 8 (C) weeks of treatment. BPaO, black circles; S587_25_PaO, light red squares; S587_50_PaO, dark red triangles; S876_6.25_PaO, light blue inverted triangles; S876_50_PaO, dark blue circles

**Table 5.** Lung CFU counts assessed during treatment and proportion of mice relapsing after treatment completion in experiment 6

| Regimen | Mean lung log<sub>10</sub> CFU count (±SD) |  | Proportion of mice relapsing after treatment for: |  |  |
| --- | --- | --- | --- | --- | --- |
|  | W-2 | D0 | W4 (+12) | W6 (+12) | W8 (+12) |
| Untreated | 4.04±0.12 | 6.58±0.22 |  |  |  |
| B<sub>25</sub>Pa<sub>100</sub>O<sub>100</sub> |  |  |  | 11/15 | 2/14 |
| S587<sub>25</sub>Pa<sub>100</sub>O<sub>100</sub> |  |  | 7/13 | 3/15 | 0/15 |
| S587<sub>50</sub>Pa<sub>100</sub>O<sub>100</sub> |  |  | 2/15 | 0/15 |  |
| S876<sub>6,25</sub>Pa<sub>100</sub>O<sub>100</sub> |  |  |  | 2/15 | 1/15 |
| S876<sub>12,5</sub>Pa<sub>100</sub>O<sub>100</sub> |  |  | 3/15 | 0/15 |  |

W-2=1 day after aerosol infection (n= mice); D0= day of treatment initiation, 2 weeks after infection. W4=treated for 4 weeks; W4+(12) = treated for 4 weeks, held for an additional 12 weeks without treatment, then sacrificed to determine the proportion with relapse, etc. From D0, 15 mice were allocated for relapse assessment at each time point indicated by proportions in the table. One mouse in the BPa W8 (+12) group died due to a gavage accident. Two mice in the S587<sub>25</sub>PaO W4(+12) group died due to cage flooding.

## DISCUSSION

One major finding of this study is that two novel DARQs currently in Phase 1 clinical trials exhibited superior sterilizing activity when substituted for bedaquiline at the same (S587) or lower (S876) doses in the BPaL regimen. These results significantly extend prior observations (3, 4) of improved bactericidal activity with these DARQs to demonstrate their treatment-shortening potential compared to bedaquiline. In a prior study, S587 exhibited greater bactericidal activity than bedaquiline in this mouse model, including against infection with an *Rv0678* (*mmpR5*) mutant with reduced susceptibility to DARQs and prevented the development of resistance in wild-type *M. tuberculosis* as well as additional pretomanid resistance in the *mmpR5* mutant (4). Similarly, S876 had superior activity compared to bedaquiline and was active *in vitro* and *in vivo* at much lower doses, e.g., 6.25 mg/kg rather than 25 mg/kg (3). These newer DARQs also appear to have a lower risk of QT interval prolongation compared to bedaquiline based on preclinical safety studies (10, 11). Together with these promising prior data, our new findings showing superior potency of the sterilizing effects of S587 and, especially, S876 support their further clinical evaluation as new drugs capable of replacing bedaquiline to shorten the treatment duration and more effectively treat and prevent infection with *mmpR5* mutants, provided they successfully pass phase 1 trials. S876 has more potent bactericidal and sterilizing activity than S587 in combination with pretomanid and TBI-223. Although this difference was suggested in our prior studies, experiments 5 and 6 provided the first head-to-head comparisons of these next-generation DARQs in combination with pretomanid and an oxazolidinone in mice. The superior *in vivo* potency of S876 appears to be largely attributable to its more potent antibacterial activity, as the steady state plasma AUC_0.24h_ values for S876 and its major microbiologically active M3 metabolite (4.0 and 15.6 μg-h/ml, respectively) are significantly lower than those for S587 and its M3 metabolite (28.8 and 41.6 μg-h/ml, respectively) and for bedaquiline and its M2 metabolite (9.6 and 55.1 μg-h/ml, respectively) after 7 days of oral dosing (at 6.25 mg/kg for S876, 25 mg/kg for S587 and bedaquiline) in mice.

A second major finding of this study is that the novel oxazolidinone TBI-223 may replace linezolid without significant loss of sterilizing efficacy in regimens containing a DARQ and pretomanid. Linezolid’s dose- and duration-dependent hematological and neurological toxicity limits its utility for treating MDR-TB and, especially, drug-susceptible TB. TBI-223 has a superior preclinical safety profile compared to linezolid and is now being evaluated in phase 1 clinical trials. Although it is less potent than linezolid against *M. tuberculosis in vitro*, its larger therapeutic window suggested by these preclinical studies may enable comparable clinical efficacy with superior safety, making regimens based on BPa plus an oxazolidinone suitable for treatment of both MDR and drug-susceptible TB. At the 100 mg/kg daily dose tested in these experiments, the single dose mean plasma AUC_0-24h_ values for linezolid and TBI-223 are 131 and 179 μg-h/ml, respectively (12). In experiment 3 (Table 2), the addition of TBI-223 to BPa significantly increased the bactericidal and sterilizing activity of the regimen. Treatment with BPaO for 4 or 8 weeks resulted in a reduction of lung CFU that was approximately 0.5 log_10_ smaller than that observed after treatment with BPaL. However, relapse rates after 8 weeks of treatment were not different between the two regimens and after 12 weeks of treatment, there were numerically fewer relapses in mice treated with BPaO, although the difference was not statistically significant. In experiment 4 (Table 3), similar results were observed with TBI-223 and linezolid in combination with the S587Pa backbone. There were again approximately 0.5 log_10_ more CFU in the S587PaO arm compared to the S587PaL arm, a difference that was not statistically significant. After 8 weeks of treatment, there were numerically fewer relapsing mice in the S587PaL group compared to the S587PaO group, but the difference was again not statistically significant. From these experiments, we conclude that TBI-223 may be an efficacious substitute for linezolid in combinations with a DARQ and pretomanid.

Judging by the similar efficacy of S587PaO and BPaL in experiment 4, the dual substitutions of a S587 for bedaquiline and TBI-223 for linezolid may result in a regimen with at least similar efficacy compared to BPaL. Since 6-month durations of BPaL have successfully treated approximately 90% of patients with XDR-TB and treatment-refractory MDR-TB (1, 2, 13, 14) and both S587 and TBI-223 have demonstrated potential safety advantages over bedaquiline and linezolid, respectively, in preclinical toxicity studies, S587PaO may allow an extended spectrum of clinical use that includes drug-susceptible TB without the dose- and duration-dependent toxicity of linezolid and with fewer concerns about QTc prolongation by bedaquiline.

The third major finding is that sutezolid, another oxazolidinone now in phase 2 clinical trials, had superior bactericidal and sterilizing activity compared to linezolid when combined with S587 and pretomanid. These results suggest that regimens combining sutezolid, which may also have lower potential for mitochondrial toxicity than linezolid (8), with a next-generation DARQ and pretomanid could result in regimens superior to BPaL in both safety and efficacy.

In summary, we present evidence from a well-established mouse model of TB that replacement of bedaquiline with safer and more effective diarylquinolines (e.g., S587 or S876) and replacement of linezolid with safer and at least as efficacious oxazolidinones (e.g., TBI-223 or sutezolid) in the clinically successful BPaL regimen may lead to superior regimens capable of treating both drug-susceptible and drug-resistant TB more effectively and safely.

## MATERIALS AND METHODS

### PK analysis

Single-dose and multi-dose plasma PK studies were carried out by BioDuro Inc. (Beijing, China). S587, formulated as described above, was administered by oral gavage at 12.5 and 25 mg/kg to uninfected BALB/c mice (n=3) for 7 consecutive days. Blood was obtained on day 1 at 1, 2, 4, 8, and 24 hrs after administration and on day 7 at 1, 2, 4, 8, 24, 48, 72, and 96 hrs after administration. Parallel analyses were carried out for S876, administered at 3.125 and 6.25 mg/kg, and for bedaquiline at 12.5 and 25 mg/kg. Plasma samples were subjected to liquid chromatography-tandem mass spectrometry using the API 4000 platform (AB Sciex, USA) for quantification of the antibiotic of interest using multiple reaction monitoring. PK parameters, including AUCs, T_1/2_, T_max_ and maximum drug concentrations (Cmax) were determined by non-compartmental analysis using Phoenix WinNonLin PK software v6.4 (Certara, USA).

### Bacterial strain

*M. tuberculosis* H37Rv was used to infect mice in these studies. The MICs of bedaquiline, S587, S876 and pretomanid against this strain were previously described (3, 4). The MICs of linezolid and TBI-223 were determined head-to-head in 3 separate experiments using the broth macrodilution method in complete 7H9 media without Tween 80 and doubling dilutions of each drug. The concentration range tested was: 0.25 – 64 μg/ml. The geometric mean MICs of linezolid and TBI-223 were 1 and 3.175 μg/ml, respectively.

### Infection model

All animal procedures were conducted according to relevant national and international guidelines and approved by the Johns Hopkins University Animal Care and Use Committee. Female BALB/c mice, 6 weeks old, were aerosol-infected with approximately 4 log_10_ CFU of *M. tuberculosis.* Treatment started 2 weeks later (D0). Mice were sacrificed for lung CFU counts on the day after infection and D0 to determine the number of CFU implanted and the number present at the start of treatment, respectively.

### Antibiotic treatment

Mice were randomized to different treatment groups. Bedaquiline was administered in all experiments at 25 mg/kg. Depending on the experiment, S587 was administered at 5, 10, 12.5, 25, 50, or 100 mg/kg, as indicated in a subscript in the results tables and graphs. S876 was administered at 6.25 mg/kg, except that a 12.5 mg/kg dose arm was also included in experiment 6. Pretomanid, linezolid, and TBI-223 were dosed at 100 mg/kg in all experiments. Sutezolid was dosed at 50 mg/kg. BDQ, S587 and S876 were formulated in 20% hydroxypropyl-β-cyclodextrin solution acidified with 1.5% 1N HCl. Pretomanid was prepared in the CM-2 formulation as previously described (15). Linezolid, sutezolid, and TBI-223 were prepared in 0.5% methylcellulose. Drugs were administered once daily by gavage, 5 days per week. Pretomanid was administered together with the diarylquinoline, 4 hrs before an oxazolidinone was given.

### Evaluation of drug efficacy

Assessments of bactericidal activity were based on lung CFU counts after 4, 6, or 8 weeks of treatment, depending on the experiment. Assessments of sterilizing activity were made 12 weeks after the completion of different durations of treatment, as indicated in the Results. At each time point, lungs were removed aseptically and homogenized in 2.5 ml PBS. Lung homogenates were plated in serial dilutions on 0.4% charcoal-supplemented 7H11 agar supplemented with 10% oleic acid, bovine albumin, sodium chloride, dextrose and catalase (OADC) and with selective antibiotics: cycloheximide (100 μg/ml), carbenicillin (100 μg/ml), polymyxin B (400,000 U/ml), and trimethoprim (40 μg/ml). For relapse assessment, the entire lung homogenate was plated.

### Statistical analysis

Group mean CFU counts were compared by one-way ANOVA with Dunnett’s correction to control for multiple comparisons. Relapse proportions were compared by Fisher’s exact test. Nonlinear regression analysis was used to fit a 4-parameter sigmoidal dose-response curve and estimate the ED_90_ and the S587 dose that would have resulted in the same mean CFU count as BDQ in the BPaL regimen. An S587 dose of 0.1 mg/kg was substituted for zero prior to log transformation. Statistical analyses used GraphPad Prism version 9.

